# First report of the East-Central South African Genotype of Chikungunya Virus in Rio de Janeiro, Brazil

**DOI:** 10.1101/082131

**Authors:** Thiara Manuele Alves de Souza, Elzinandes Leal de Azeredo, Jéssica Badolato Corrêa da Silva, Paulo Vieira Damasco, Carla Cunha Santos, Fabienne Petitinga de Paiva, Priscila Conrado Guerra Nunes, Luciana Santos Barbosa, Márcio da Costa Cipitelli, Thaís Chouin-Carneiro, Nieli Rodrigues da Costa Faria, Rita Maria Ribeiro Nogueira, Fernanda de Bruycker-Nogueira, Flavia Barreto dos Santos

**Affiliations:** Viral Immunology Laboratory, Oswaldo Cruz Institute, Rio de Janeiro, Brazil; Medical and Surgery School, Federal University of the State of Rio de Janeiro UNIRIO, Medical Sciences Faculty, University of the State of Rio de Janeiro, UERJ; Rio Laranjeiras Hospital, Rio de Janeiro, Brazil; Flavivirus Laboratory, Oswaldo Cruz Institute, Rio de Janeiro, Brazil

**Keywords:** Chikungunya, Rio de Janeiro, Brazil

## Abstract

**Background:** Chikungunya virus (CHIKV) is an arbovirus that causes an acute febrile illness characterized by severe and debilitating arthralgia. In Brazil, the Asian and East-Central South African (ECSA) genotypes are circulating in the north and northeast of the country, respectively. In 2015, the first autochthonous cases in Rio de Janeiro, Brazil were reported but until now the circulating strains have not been characterized. Therefore, we aimed here to perform the molecular characterization and phylogenetic analysis of CHIKV strains circulating in the 2016 outbreak occurred in the municipality of Rio de Janeiro.

**Methods:** The cases analyzed in this study were collected at a private Hospital, from April 2016 to May 2016, during the chikungunya outbreak in Rio de Janeiro, Brazil. All cases were submitted to the Real Time RT-PCR for CHIKV genome detection and to anti-CHIKV IgM ELISA. Chikungunya infection was laboratorially confirmed by at least one diagnostic method and, randomly selected positive cases (*n=*10), were partially sequenced (CHIKV E1 gene) and analyzed.

**Results:** The results showed that all the samples grouped in ECSA genotype branch and the molecular characterization of the fragment did not reveal the A226V mutation in the Rio de Janeiro strains analyzed, but a K211T amino acid substitution was observed for the first time in all samples and a V156A substitution in two of ten samples.

**Conclusions:** Phylogenetic analysis and molecular characterization reveals the circulation of the ECSA genotype of CHIKV in the city of Rio de Janeiro, Brazil and two amino acids substitutions (K211T and V156A) exclusive to the CHIKV strains obtained during the 2016 epidemic, were reported.

## Introduction

Chikungunya virus (CHIKV) is an arbovirus belonging to the Togaviridae family and Alphavirus genus that causes an acute febrile syndrome with severe and debilitating arthralgia^1–4^. It belongs to the Semliki Forest Complex and the viral particle is small, composed of an icosahedral capsid surrounded by a lipid envelope that measures approximately 60-70 nm in diameter and the genome consists of a positive sense single stranded RNA measuring 12kb in length, which encodes four non-structural proteins (NSP1-4) and five structural proteins (C, E3, E2, 6K and E1) ^5–7^.

CHIKV was first described in 1952 on the Makonde Plains, along the border between Tanzania and Mozambique (East Africa), and since its discovery, has been responsible for important emerging and re-emerging epidemics in several tropical and temperate regions of the world^8^. So far, distinct genotypes of CHIKV have been identified as West African, East-Central South African (ECSA) and Asian. Besides, the Indian Ocean Lineage (IOL) has emerged in Kenya during 2004 as a descendant lineage of ECSA and caused several outbreaks in Indian Ocean Islands, India and Asia during 2005-20114,^9,10^. Although the *Aedes* (Ae.) *aegypti* mosquito has been highlighted as the main vector for the urban cycle of CHIKV11. *Ae. albopictus* also has a high vectorial competence for the virus due to an A226V mutation in the E1 gene of ECSA genotype that generated the IOL, which promotes an increase in infectivity in the midgut, dissemination to the salivary glands and transmission by this mosquito ^9,12–14^. In addition, a large number of imported and autochthonous cases of CHIKV have been reported in American, Europe and Asian countries since 2006 due to viremic travelers arising from Africa, India and Indian Ocean islands ^11,13^.

In the Americas, the first autochthonous transmission of Asian genotype was reported during 2013 in San Martin Island, in the Caribbean ^14,15^ and since that, many autochthonous cases have emerged in Caribbean, South and Central America countries, United States, Mexico, Brazil and the Andean countries ^16^. In Brazil, the first autochthonous cases of the Asian and ECSA genotypes were reported during 2014 in the North and Northeast cities of Oiapoque (Amapá State) and Feira de Santana (Bahia State), respectively ^10,17^. In 2015, 38,332 chikungunya suspected cases distributed in 696 municipalities were reported and this amount increased to 216,102 in 2016, distributed in 2,248 municipalities until 32^nd^ epidemiological week and, from those, approximately 50% were confirmed by clinical epidemiological criteria in the Northeast (Alagoas, Bahia, Ceará, Maranhão, Paraiba, Pernambuco and Rio Grande do Norte municipalities) and Southeast of the country (Rio de Janeiro and São Paulo municipalities) ^18^. Despite the Northern region of Brazil presented the highest incidence of chikungunya cases, the virus has spread to the Southeast region in 2015 and it resulted in 18,173 cases during 2016, with 13,058 of those restricted to the city of Rio de Janeiro^18^.

The exponential growth of chikungunya cases in Rio de Janeiro represents a serious public health problem, especially due to the current co-circulation with dengue and zika. As both Asian and ECSA genotypes were introduced in Brazil in 2014, the viral surveillance is of great importance to access the impact over a population, as the role of distinct genotypes in the disease severity and chronicity is not well understood. Moreover, the monitoring and characterization of CHIKV genotypes allow the identification of possible mutations such as the E1-A226V, of described epidemiological impact ^9,12^. Despite the increased incidence of the disease in the past year, the information of CHIKV genotypes circulating in Brazil is still scarce. Here, we aimed to perform the genotype characterization of CHIKV strains detected during the ongoing 2016 outbreak in Rio de Janeiro, Brazil.

## Material and Methods

### Ethical Statement

The samples analyzed in this study were from the an ongoing project for arbovirus research in Rio de Janeiro, Brazil approved by resolution number CSN196/96 from the Oswaldo Cruz Foundation Ethical Committee in Research (CEP 111/000), Ministry of Health-Brazil. All participating subjects provided a written consent.

### Clinical samples

The plasma samples analyzed in this study were collected from April 2016 to May 2016 during the chikungunya outbreak in Rio de Janeiro, Brazil. Patients were assisted at the Hospital Rio Laranjeiras (HRL) where an infectious disease physician collected data on demographic characteristics, symptoms and physical signs using a structured questionnaire. Chikungunya suspected cases (*n=*91) were obtained during an active surveillance performed by the Laboratory of Viral Immunology, IOC/FIOCRUZ. All cases were submitted to the Real Time RT-PCR for CHIKV genome detection^19^ and to the anti-CHIKV ELISA IgM kit (Euroimmun, Lubeck, Germany), according to the manufacturer’s protocol. Chikungunya infection was laboratorially confirmed by at least one diagnostic method in 76.97% (70/91) of the cases, 48.57 (34/70) by serology and 84.28 (59/70), by Real Time RT-PCR. Moreover, 35.85% (23/70) of the cases were confirmed by both methods. Chikungunya positive cases (*n=*10) by Real Time RT-PCR, were randomly selected for partial sequencing (E1 gene) and phylogenetic analysis.

### Chikungunya virus genome amplification, sequencing and phylogenetic analysis

Viral RNA was extracted from 140μL of plasma samples using QIAamp Viral RNA Mini Kit (Qiagen Inc., Germany) according to the protocol described by the manufacturer and stored at −70°C.

For partial sequencing of CHIKV E1 gene, primers by^20^ were used in two steps with Qiagen OneStep RT-PCR Kit (Qiagen, Inc., Germany). Five microliters of the extracted RNA was reverse transcribed into cDNA and amplified using sense _102465_′-TTACCCNTTYATGTGGGG-3′_10262_ and antisense _107935_′-CTTACSGGGTTTGTYGC-3′_10777_ primers, on thermocycling parameters of one cycle of reverse transcription (50°C/60 min), one cycle for activation of hotstart polymerase enzyme, reverse transcriptase inactivation and degradation of the template for the cDNA (95°C/15min) followed by 40 cycles of denaturation (94°C/30 sec), annealing (60°C/3 min) and extension (68°C/30 sec), ending with a final extension cycle (68°C/10 min), in a GeneAmp^®^ PCR System 9700 (Applied Biosystems^®^, California, USA). The volume of 0,5μL of PCR products were submitted to semi-nested PCR using primers sense in combination with primer _107145_′-TRAAGCCAGATGGTGCC-3′_10698_ for amplification of a 469 bp fragment, on thermocycling parameters of one cycle of denaturation (94ºC/2 min), followed by 40 cycles of denaturation (95°C/30 sec), annealing (55°C/1 min) and extension (72°C/30 sec), ending with a final extension cycle (72°C/5 min), also in a GeneAmp^®^ PCR System 9700 (Applied Biosystems^®^, California, USA).

The fragments generated were purified using PCR Purification Kit or Gel Extraction Kit (QIAGEN, Inc., Germany) and sequenced in both directions using the BigDye Terminator Cycle Sequencing Ready Reaction version 3.1 kit (Applied Biosystems^®^, California, USA). The thermocycling conditions consisted of 40 cycles of denaturation (94°C/10 sec), annealing (50°C/5 sec) and extension (60°C/4 min). Sequencing was performed on an ABI 3730 DNA Analyzer, Applied Biosystems^®^, California, USA^21^.

The sequences analysis was performed using BioEdit (http://www.mbio.ncsu.edu/bioedit/bioedit.htmL), sequences’ identity was performed using BLAST (http://blast.ncbi.nlm.nih.gov/Blast.cgi) and alignments using CLUSTAL OMEGA (http://www.ebi.ac.uk/Tools/msa/clustalo/). Phylogenetic trees were constructed using the MEGA 6 (http://www.megasoftware.net/), by the "Neighbor-Joining" method, Kimura-2 parameter model (K2), with a bootstrap of 1,000 replications. The tree was built based on the analysis of the best fit for model, as provided by the software. Partial CHIKV genome sequences were deposited in GenBank and accession number were as follow: KX966400 to KX966409.

## Results

The molecular characterization and phylogenetic analysis of representative strains (*n=*10) of CHIKV detected in infected patients during the 2016 outbreak in Rio de Janeiro was performed in comparison to reference sequences available on Genbank and were used to represent the Asian, ECSA and West Africa genotypes. The results showed that all the analyzed strains grouped in the ECSA genotype branch, together with a sequence from a sample identified in Bahia in 2014 (Figure). Additionally, the molecular characterization of the E1 fragment revealed that the alanine amino acid was present at the E_226_ position, showing no A226V mutation. Interestingly, a K211T amino acid substitution was identified in all analyzed samples and a V156A substitution was identified in two samples of this study. Furthermore, the CHIKV sequences identified in Bahia belonging to the ECSA genotype did not show this substitution at the 211 amino acid and where a lysine (K) is found in the prototype.

**Figure:**
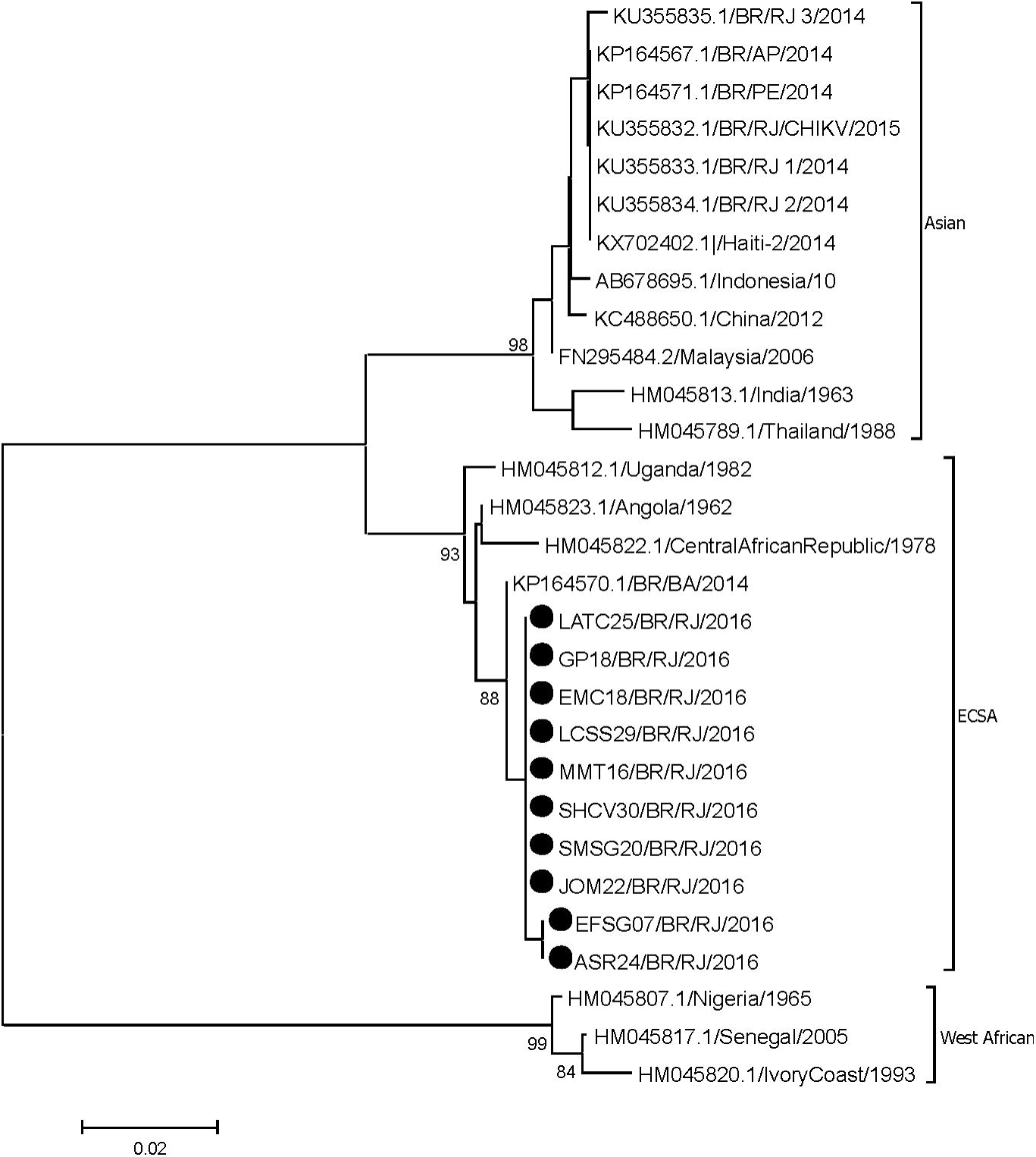
Genotyping of CHIKV strains (*n=*10) identified in Rio de Janeiro during the outbreak occurred in 2016. Neighbor Joining method, K2 parameters model, bootstrap of 1,000 replications. The CHIKV sequences analyzed are represented by black circles. CHIKV strains were named as follows: GenBank accession number (or name strain)/country/year.

## Discussion and Conclusions

Since its first description, CHIKV has been responsible for important emerging and reemerging epidemics of arbovirus disease characterized by severe and incapacitating polyarthralgia syndrome^8,22^. Because of the intense movement of viremic travelers arising from Africa, India and Indian Ocean islands, many imported cases of this disease were reported in American, European and Asian countries since 2006 ^11,13^.

Due to the high vector density, the presence of susceptible individuals and the intense movement of people, Brazil has a major risk for the occurrence of arbovirus epidemics. After its introduction, CHIKV has caused outbreaks in many regions of Brazil, mainly affecting the Northern region. Despite that, the Southeast Region has played an important role in the disease epidemiology as imported cases are reported since 2010, and where stands out with the most cases of autochthonous transmission of this virus during 2015 and 2016 ^23,18^.

The exponential growth of CHIKV cases in Rio de Janeiro represents a serious public health problem and the co-circulation of three arboviruses (DENV, CHIKV and ZIKV) results in difficult differential diagnosis of those diseases ^24^. Prior to this study, no phylogenetic information was available on the autochthonous CHIKV strains circulating in Rio de Janeiro and, the data available was from the Asian genotype characterized in imported cases analyzed in 2014 and 2015 ^25^.

From our knowledge, this is the first report on the ECSA genotype circulation during the 2016 outbreak in Rio de Janeiro. This genotype was first reported in Feira de Santana, Bahia, Northeast region of Brazil, during 2014 and studies revealed that the strains originated from Angola (West Africa). Moreover, it was the first time that this genotype was reported in Americas. The other CHIKV introduction in Brazil was from the Asian genotype in Oiapoque, Amapá, North Brazil, also during 2014 and, studies revealed those strains were originated from the Caribbean and South that America^10,17,26^ Additionally, the molecular characterization the E1 gene fragment analyzed showed that an alanine was present at the E226 position, therefore showing no A226V mutation. Studies performed during the 2005-2006 epidemic occurred in the Reunion Island characterized that this mutation was responsible for generating the IOL, responsible for an increased CHIKV transmission by the vector Ae. Albopictus ^9,12–14^. Furthermore, the E1 gene represents a target region for this analysis due to the high antigenic variability, role in the attachment, viral entry into target cells and viral replication during CHIKV infection ^7,27^. However, this study revealed a K211T amino acid substitution in all samples analyzed and a V156A substitution in two sequences. The former substitution was not identified in the strains from Bahia, which has a Lysine (K) as the prototype of this genotype. Further studies are needed to clarify the consequences of those mutations, including to the mosquitoes fitness and the human immune system, but other studies suggest that new mutations such as L210Q, I211T and G60D in the E2 region of the IOL also offer advantages for the transmission of CHIKV by *Ae. Albopictus* ^12,28,29^. The mutations K211E on E1 and V264A on E2 were reported to impact *Ae. aegypti* 's fitness in India during the 2006 to 2010 epidemic ^30,31^.

This study provides the first genotype surveillance of autochthonous CHIKV cases during the 2016 epidemic in Rio de Janeiro and stress the need for monitoring the spread of the distinct genotypes and the identification of possible mutations that may facilitate the viral transmission by the mosquitoes’ vectors. Notwithstanding, none of the chikungunya patients were hospitalized or had other complications not related to classic rheumatologic chikungunya syndrome. Rio de Janeiro is an important port of entrance and spread of arboviruses, as observed for the distinct DENV serotypes, especially due to its high vector density, susceptible individuals and intense tourists movement. The recent events occurred in Rio de Janeiro also reinforces the need for viruses’ surveillance and characterization.

## Authors’ contributions

FBS and FBN designed the study. TMAS and FBN implemented the sequencing study, analyzed the data and wrote the paper. PCGN, JBCS, FPP, LSB and MCC collected and processed the samples. TCC and NRCF analyzed the data. PVD and CCS assisted the patients during cases investigation and samples collection. ELA and RMRN provided the laboratory structure and funding for carrying out the experiments. FBS is the guarantor of the paper.

## Equal contribution

Flavia Barreto dos Santos and Fernanda de Bruycker-Nogueira contributed equally to the work.

## Acknowledgements

To Dr Ana Maria Bispo de Filippis, Head of the Flavivirus Laboratory, IOC/FIOCRUZ for lab support, to the staff of the Rio Laranjeiras Hospital and to the Parasitary Biology Postgraduate Program at Oswaldo Cruz Institute/FIOCRUZ.

